# Optofluidic Fiber Component for Separation and counting of Micron-Sized Particles

**DOI:** 10.1101/2021.04.13.439593

**Authors:** T. Kumar, A.V. Harish, S. Etcheverry, W. Margulis, F. Laurell, A. Russom

## Abstract

An all-fiber separation component capable of sorting and counting micron-sized particles based on size is presented. A sequence of silica fiber capillaries with various diameters and longitudinal cavities were used to fabricate the component for separation and detection in an uninterrupted flow. Fluorescence microparticles of 1 μm and 10 μm sizes are mixed in a visco-elastic fluid and infused into the all-fiber separation component. Elasto-inertial forces focus the larger particle to the center of the silica capillary, while the smaller microparticles exit from a side capillary. Analysis of the separated particles at the output showed a separation efficiency of 100% for the 10 μm and 97% for the 1 μm particles. In addition, the counting of the larger particles is demonstrated in the same flow. The separated 10 μm particles are further routed through another all-fiber component for counting. A counting speed of ~1400 particles/min and with the variation in amplitude of 10% is achived. A combination of separation and counting can be powerful tool may find several applications in biology and medicine, such as separation and analysis of exosomes, bacteria, and blood cell sub-populations.

## 1. Introduction

Separation of cells, bacteria, and organic debris from a mixed population is important in life-sciences, especially in cytology [1][2]. One prime example is the need to detect a few circulating tumour cells in the blood, which are the source of metastasis and carry information about the origin of cancer [3]. Separation of bacteria from blood cells is also useful for early diagnostics of blood stream infections and for identification of possible treatment with antibiotics. Microfluidics is an emerging technology for high efficiency and high throughput cell separation [4] [5][6]. Microfluidics technology allows for the fabrication of miniature devices with very low cost, opening the possibility of point-of-care (POC) diagnosis (i. e. where the patient is) [7]. Progress in microfluidic and POC diagnosis is expected to be game-changer technology, particularly in developing countries with limited access to medical facilities [8]. Microfluid-based cell separation has been demonstrated using affinity biomarkers, with active techniques such as magnetic and acoustic separation [9], and by passive techniques such as inertial [7] and elasto-inertial [10] microfluidics. The latter two have recently gained attention due to their simplicity, high throughput, the capability of processing huge volumes of sample, and low stress on cells. Inertial microfluidics relies on inertial forces that are acting on the flowing particles or cells at high flow-rate through a microfluidic channel. Depending on the geometry of the channel, these forces drive the particles to a specific streamline position, allowing for particle focusing or separation from a mixed population. Elasto-inertial microfluidics combines the inertial forces with additional viscoelastic forces that occur when using non-Newtonian fluids [11]. This allows extending the applicability of inertial microfluidics by covering a larger range of particle sizes and channel geometries. Significant progress has been made in microfluidic-based cell separation [12][13]. However, a miniaturized device for cell analysis in a POC setting should include, besides cell separation, the capability of detecting and counting the cells [14]. To date, microfluidic devices for cell separation mostly rely on bulky flow-cytometers or external microscopes for detection, which increases the overall size and cost of the devices and prevents its use in POC settings. On the other hand, integrated and miniaturized flow cytometers have been demonstrated by using waveguides or embedded optical fibers [15][16] but they lack the capability of separating the cells of interest from the sample.

In our previous work, we have used hollow optical fibres to detect, trap, collect and analys microparticles in fluids [17][18]. The system was further develop to a compact all-fiber flow cytometer by assembling silica optical fibers and microcapillaries [19]. It used elasto-inertial microfluidics in the capillaries for particle focusing and an optical fiber for carrying the light to and from the particles. This was a high-performance device fabricated at a low cost without the need for clean-room facilities.

Here, we present a silica-based component that allows for the separation of particles of different sizes with high efficiency. In our proof-of-principle experiment, a sample consisting of a mixture of 10 μm and 1 μm particles is flown into the all-fiber separation component, where particles of different sizes migrate away from each other and follow independent outlets. The results show that 100% of the 10 μm particles exit the separation chamber through a central outlet, while 97% of the 1 μm particles exit through a side outlet (and 3% in the central outlet). Finally, an experiment is performed to demonstrate that integration of size-dependent separation and fluorescence-based detection of particles is possible. After separation, the larger particles are counted in a miniaturized all-fiber cytometer in continuous flow. The present work shows that fiber technology provides a promising platform to build modular, flexible and efficent optofluidic systems for various biological applications.

## 2. Experimental Methods

### 2.1. Experimental setup

The experimental setup for separation of particles based on size is shown in Fig. 1. The all-fiber separation component has two inlets, one for feeding pure viscoelastic fluid (sheath) and the other carrying the sample containing the microparticles. Two microfluidic pumps are used to push the sheath and the sample into the separation component. In the current experiment, fluorescent microsphere suspensions with 10 μm (green) and 1 μm (red) were used [20].

**Figure: 1.**
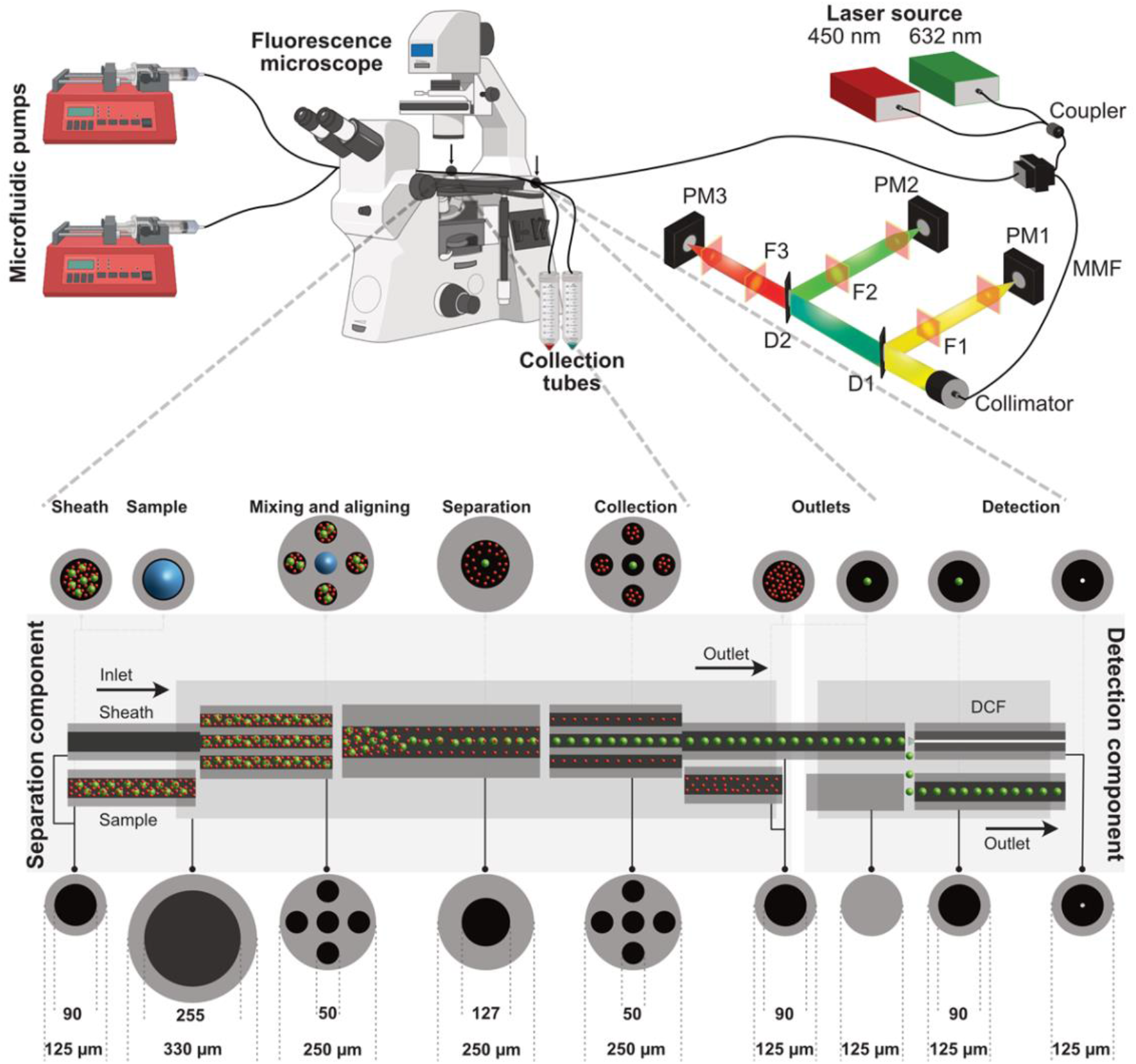
Experimental setup for separating the 10 μm (green) and 1 μm (red) fluorescent microparticles using an all-fiber separation component. The microfluidic pumps push PEO solution and PEO solution mixed with fluorescent microparticles. The separated particles exit the component in two distinct capillaries and the 10 μm fluorescent microparticles are counted. Inset crossection show the dimension of the silica fiber components at each section of the components.

To characterize particle focusing, a fluorescence microscope is used to image different sections of the all-fiber component. When in operation, the fluorescence microscope can be used to image the streamlines of the fluorescent particles inside the fiber capillary to distinguish the particles. On the detector side a Fluorescein isothiocyanate (FITC) filter cube was used for the 10 μm fluorescent particles with excitation/emission peaks of 468/508 nm (Green) and a Tetramethylrhodamine (TRITC) filter cube for the 1 μm fluorescent particles with excitation/emission peaks of 542/612 nm (Red). The images were processed and analyzed using the software ImageJ (NIH, MD, USA). Fluorescent microscope images were taken using a motorized Nikon Ti-Eclipse inverted microscope coupled with a Lumencor SOLA light engine as the excitation source. The separation component has two outlets, which are collected in two different Eppendorf for further analysis, or flow into additional all-fiber devices, such as a flow-cytometer.

The sheath and sample are pushed into the all-fiber separation component using a mid-pressure syringe pump neMESYS, Cetoni. Two syringes of 25 mL are used to pump the fluids into the separation component. The flowrates for both the sheath and the sample are controlled independently using neMESYS user interface software.

### 2.2. Design and fabrication of all-fiber separation component

The schematic diagram of the separation component is shown in Fig. 1. It uses an assembly of four different silica fiber capillaries, the cross-sections of which are shown in Fig. 1. All inlets and outlets consist of capillaries with 90 μm inner diameter and 125 μm outer diameter (90/125 μm). Separation takes place in a 127/250 μm separation channel, where the mixture of microparticles flow with the viscoelastic fluid, as shown in Fig. 1. The length of the separation channel is 3 cm. The input mixture is fed to the separation channel through a short piece of 5-hole fiber of length less than 1 mm. The central hole of 5-hole fiber carries only the sheath and the outer four holes carry the mixture of microparticles. This is accomplished by splicing one of the 90/125 μm capillaries to the central hole of the 5-hole fiber leaving room for the mixture to enter from the outer holes. Part of the structure is encased in a housing capillary with dimensions 250/330 μm. At the output, a similar arrangement with 5-hole fiber with central hole spliced to 90/125 μm capillary is made to collect the separated microparticles. This 5-hole fiber guarantees that the central collection capillary is aligned with the large focused particles and the four outer holes provide low flow resistance for the collection of the smaller particles. The distance between the separation channel and the 5-hole fiber is fixed at 40 μm so that the particles do not deviate significantly from the separated path. UV-curing glue is used to seal the housing capillary at both ends, preventing leakage.

After separation by size, the identification by fluorescence and counting of the 10 μm microparticles is carried out. An all-fiber cytometer like the one reported in Ref. [19] was built and integrated to the separation component as shown in Fig 1 labelled detection part. The all-fiber cytometer has a 56/125 μm input capillary where particle focusing takes place and a 90/125 μm output capillary to remove particles after counting. It is fabricated by aligning a double-clad fiber to the input capillary. This fiber provides laser excitation from the 9 μm central core, and collects back-scattering and fluorescence in the coaxial outer core. Besides the input and output capillaries and the double-clad fiber, the device incorporates a piece of dummy fiber used for simplifying alignment. An external 250/330 μm housing capillary holds the assembly together as shown in Fig 1.

### 2.3. Elasto-Inertial microfluidic setup

The viscoelastic (non-Newtonian) fluid used in this work was prepared using the polymer polyethylene oxide (PEO) with an average molecular weight (Mv) of 2000,000 (Sigma-Aldrich). Powdered PEO was weighed and added directly to a freshly prepared 1X Phosphate buffer saline (PBS) solution. The mixture was left in a magnetic stirrer overnight for mixing. The concentration of PEO used in the experiment was 500 PPM. The flow of particles was tracked and observed under a fluorescence microscope.

## 3. Results and Discussion

Various separation channel length and flow rates of the fluids into the separation component were investigated. As expected, the separation efficiency varied depending on the length of the separation channel and flow rates of sheath and sample. Figure 2A shows our investigation of the length of the separation channel for separation efficiency. The flow rate of the sample and sheath is kept constant at 10 μL/min. Once the sample and sheath enters the separation channel the separation of the 10 and 1 micron particles is monitored at different positions of 1, 2, 3, 4 and 5 cm from the entrance as shown in Fig. 2A. Images were taken at these positions by shining blue excitation light to illuminate the green fluorescent 10 μm microparticles and then green light to illuminate the 1 μm red fluorescent particles. At a distance of 3 cm from the entrance to the separation channel, distinct red and green streamlines are seen which signifies the separation has occurred. Hence a separation channel length of 3 cm was used in our separation component.

**Figure: 2.**
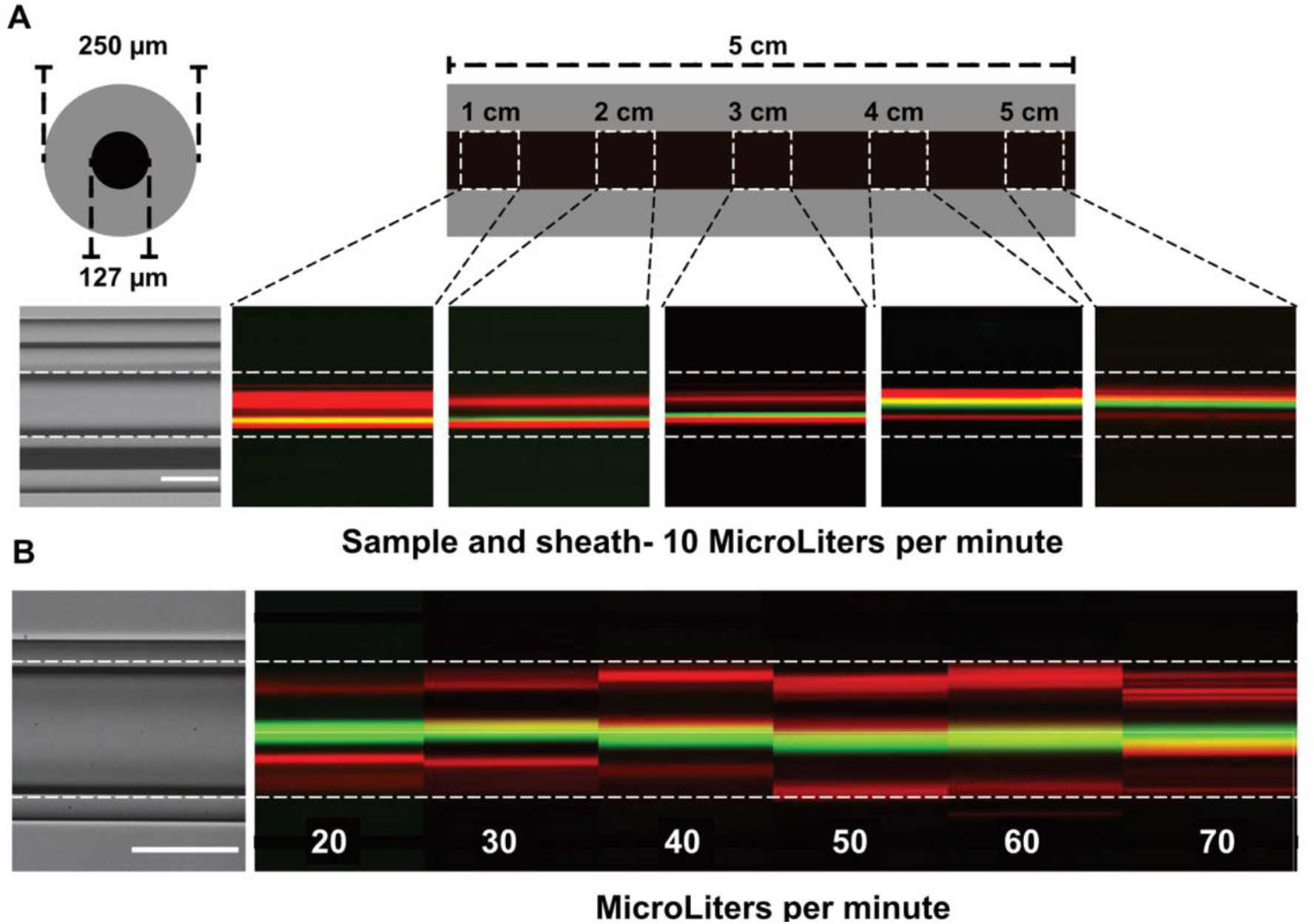
Experimental results showing the separation efficiency at A) different lengths of the separation component, i.e., 1, 2, 3, 4 and 5 cm from the entrance of the separation component, and B) different flow rates of sheath from 10 to 70 μL/min and sample kept constant. The red lines in the figure indicate the streamlines of the 1 μm red fluorescent particles, the green lines are the steamlines for 10 μm green fluorescent particles. Yellow lines appear when the green and red streamlines overlap.

Additionally, the flow rate of sample to the separation component was also investigated. It was varied from 10 to 70 μL/min while keeping the sheath constant at 10 μL/min. The streamline of green and red microparticles were monitored at 3 cm from the entrance to the separation channel as shown in Fig. 2B. At 10 μL/min of the sample flow rate the green particles focused at the center in a narrow streamline (Fig. 2A). For a sample flow rate of 20 μL/min the green particles are separated from red but there is no strong focusing. For higher flow rates the separation degrades further as seen in Fig. 2B. Finally, we varied the sheath flow rate from 10 to 70 μL/min keeping the sample flow rate at 10 μL/min. The best separation was observed with a sample flow rate of 10 μL/min and a sheath flow rate of 40 μL/min for a 3 cm long separation channel.

The results of the experiment with the separation component are shown in Fig. 3. Fig. 3a shows a brightfield (BF) image of the separation component at the end of the separation channel. After exiting the separation channel the particles are spatially separated. All the 10 μm green fluorescent particles enter the central hole of the 5-hole capillary as seen in Fig. 3c, while the 1 μm, red fluorescent particles enter the outer four holes, as seen in Fig. 3b. In order to understand the particle distribution at the end of the separation capillary we analyse the images taken with the microscope. Using Fig. 3b and 3c the intensity of the light along the diamter vs. the diameter of the separation capillary were plotted in Fig. 3d. For the 10 μm green fluorescent plot there is only one central peak appearing at the center of the migration capillary. This shows that all the 10 μm green fluorescent particles are focused in the center of the separation capillary. However, in the plot for 1 μm red fluorescent particles there is a large central peak and two smaller peaks on either side. The central big peak in case of 1 μm red fluorescent particle consist of a superposition of light from two output capillaries projected on top of each other in the image. To check the separation efficiency, the particles collected at the outlets of the component is further analysed. The collected particles were spun and 10 microliter of the suspension was taken and loaded onto a hemocytometer. Figure 4A shows the fluorescent image taken under a microscope of the mixture of the 10 and 1 μm microparticles that was fed into the separation component. Figure 4B shows the fluorescent image of the separated samples collected at the center hole of the 5-hole capillary that primarily contains all the 10 μm green particles and very few 1 μm red particles. Figure 4C shows the fluorescent image of the sample collected from the outer 4 holes of the 5-hole fiber with red particles only. Taking pictures of 5 sets of 16 squares (4 in the corner and 1 in the center) manual counting was performed in all the squares and the results were calculated for both 1 and 10 μm particles. The results showed 100% separation of 10 μm particles and 97% of 1 μm particles as can be seen in the bar chart of Fig. 4D.

**Figure: 3.**
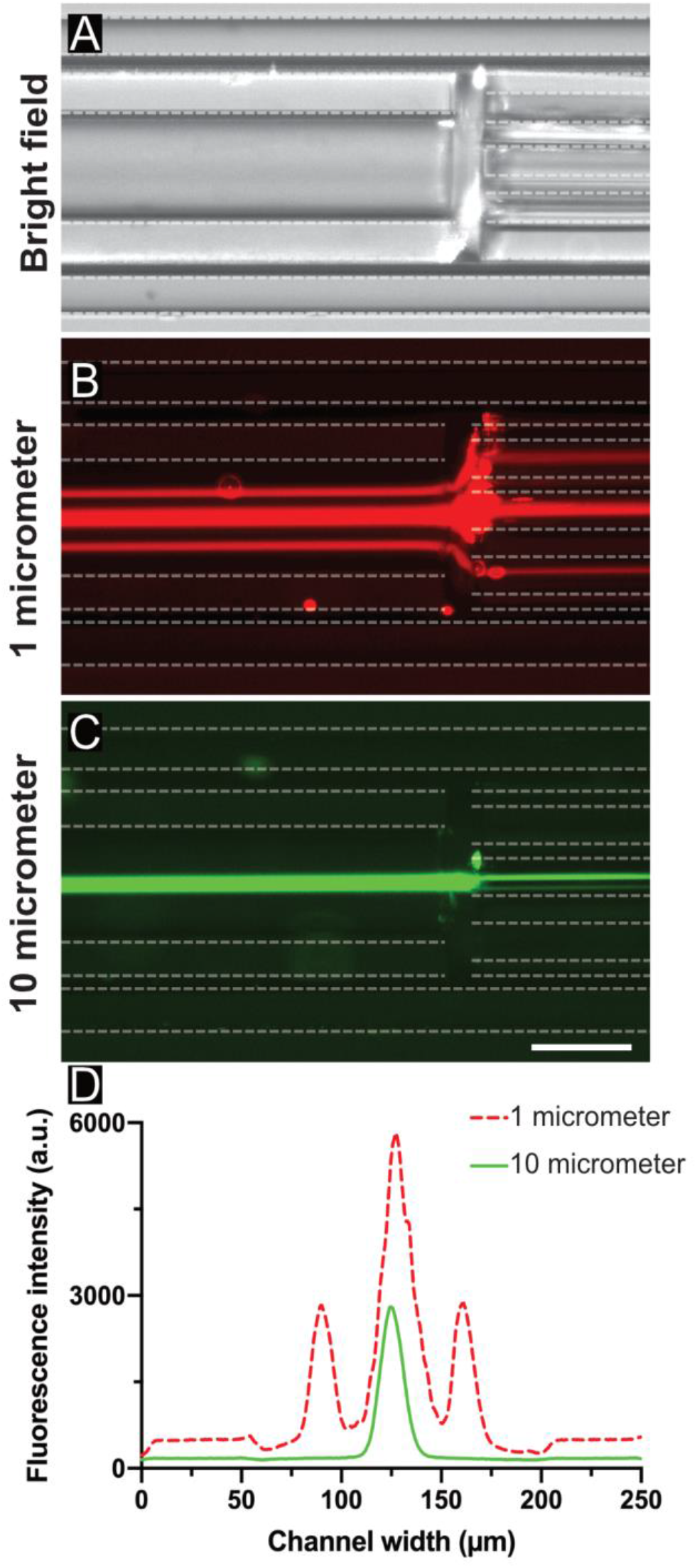
a) Brightfield image of the output of the separation channel showing the 127/250 μm capillary on the left and the 5-hole fiber on the right. b) Microscope image of the same section taken during separation where the 1 μm red particles are separated out from the center and entering the central hole of the 5-hole fiber. c) Microscope image of the same where the 10 μm green fluorescent particles are focused to the central hole of the output fiber. d) Normalized intensity graph showing focusing of the 10 μm particles (green) while the 1 μm particles (red) were separated.

**Figure: 4.**
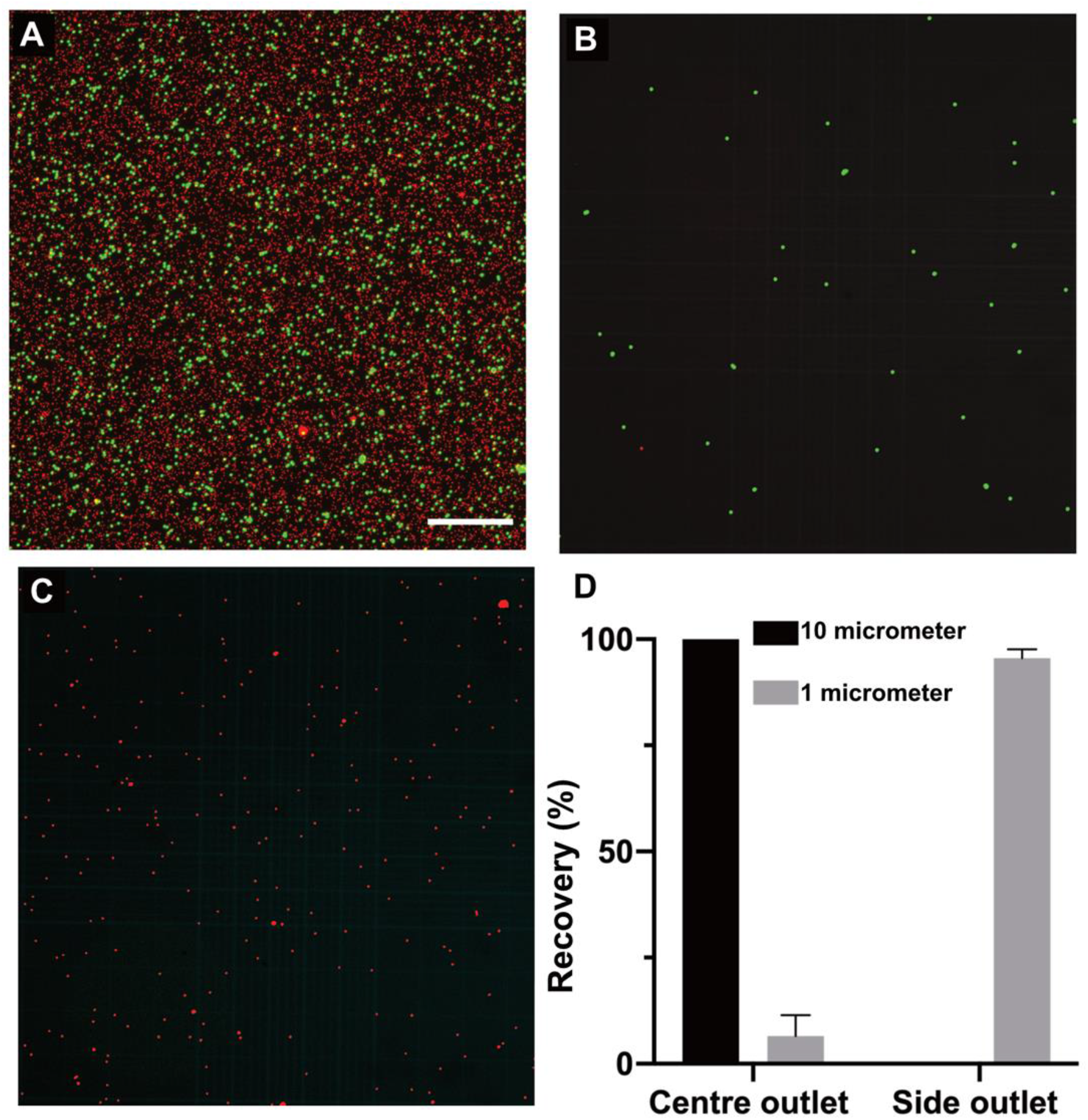
Enrichment analysis of separated particles. Results showing the separation efficiency of 1 and 10 μm particles calculated using a hemocytometer. A) Fluorescent images of 1 (red) and 10 (green) μm particles before processing through the separation device. B and C) Fluorescent images after processing the sample through the separation device. D) The graph shows a 100% separation of 10 μm particle and 97% of 1 μm particles.

After separation of 10 μm green fluorescent particles the particles are counted using a fiber optical partical counting unit that is connected to the separation component as shown in Fig. 1 [19]. A blue laser at 450 nm is used to illuminate the incoming fluorescent microparticles with the light guided in the inner core of a double-clad fiber. Fluorescence from the particles is collected in the inner cladding of the double-clad fiber and guided to the detector end of the fiberwhere the signal is filtered through a dichroic filter that only allows green fluorescence light to pass through. The green light is detected using a silicon photomultiplier (SiPM) and the temporal trace collected using an oscilloscope and analysed in a PC. The separated 10 μm green microparticles are counted using this all-fiber component before they are collected in an Eppendorf. After the collection of fluorescent particles from the outlets in the Eppendorfs, the 10 μm microparticles were also counted using the Bright-Line hemacytometer (Sigma-Aldrich).

Figure 5 shows the time domain trace for a time window of 30 s of the fluorescence signals collected from the all-fiber flow cytometer device. Each pulse in the time trace of Fig. 5 corresponds to one fluorescent 10 μm particle. We count each pulse above a threshold of 0.1 V and get 716 particles in 30 s. The number of particles counted per second counted depends on the flow rate of the sample. The measurement in Fig. 5a is taken with an flow rate of 10 μL/min for the sheath (PEO) and 40 μL/min for the sample. The coefficient of variation (CV) in the amplitudes of the fluorescence signals is 10%. To verify the number of microparticles counted using the all-fiber component we collected them in an eppendorf for three minutes. A coulter counter was then used to count the number of microparticles in Eppendorf. We did four sets of measurements with the coulter counter giving an average of 4100 particles in three minutes. The results are plotted in Fig. 5b. With the all-fiber component the average count was 1400 particles/min. This shows the efficacy of our all-fiber component.

**Figure: 5.**
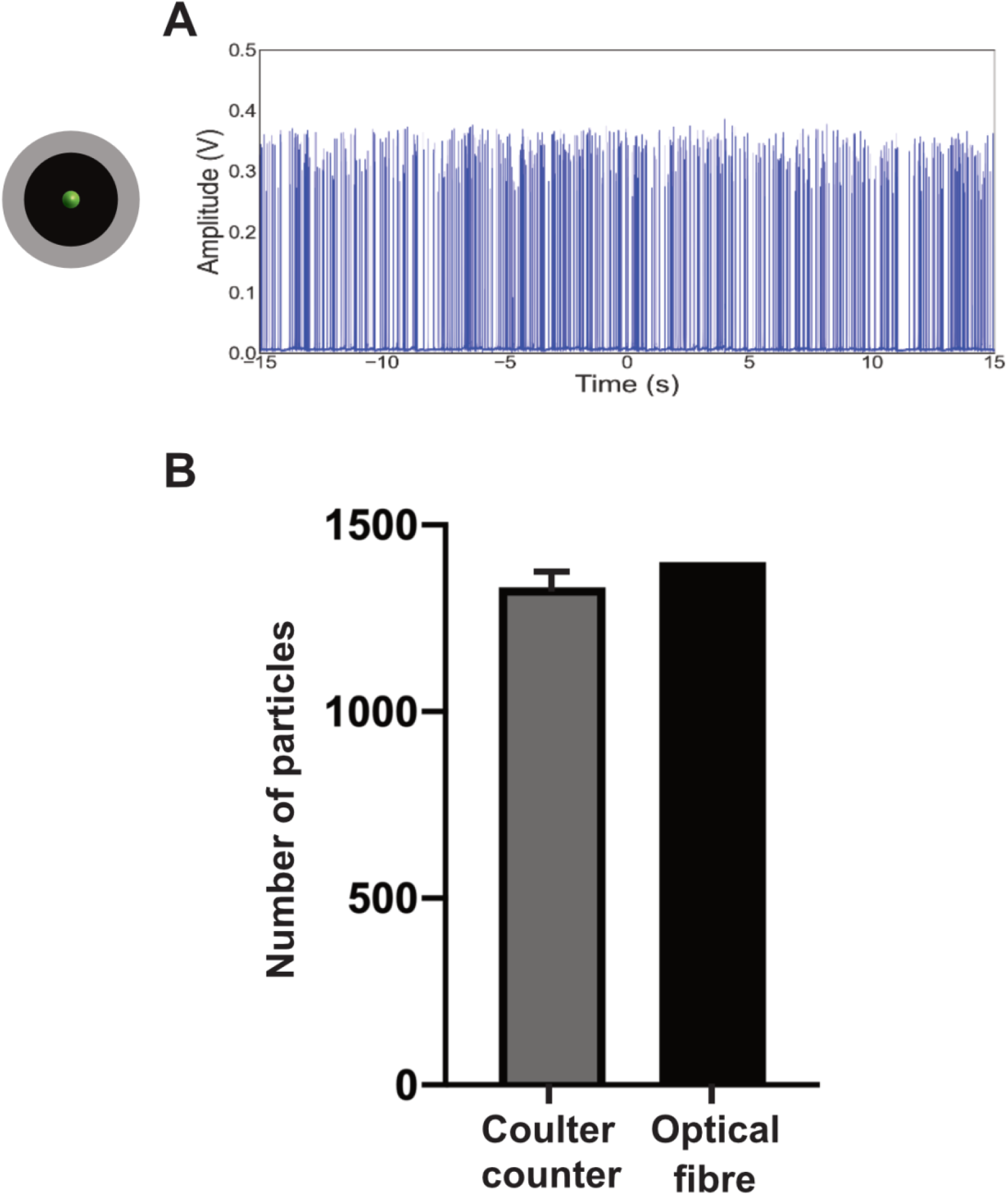
a) Temporal trace showing the pulse train where each pulse corresponds to one 10 μm green fluorescent particle counted in an all-fiber cytometer device after separation from a mixture of 10 and 1 μm particles in an all-fiber separation component and, b) a bar chart comparing the number of microparticles counted with Coulter counter and fiber component

The rates of flow to the integrated all-fiber component are mainly decided by the separation component. However, the microflow cytometer also needs specific flow rates for the 10 μm particles to focus at the center of the 90/125 μm fiber capillary. This leads to a somewhat non-uniform amplitude of the fluorescence signals as seen in Fig. 6 due to focusing fluctuations of the 10 μm microparticles. An optimized version of the device integrating separation and flow cytometry could make use of capillary diameters matched to the flow rates needed. However, such an optimization falls beyond the scope of the current work in this paper.

## 4. Conclusions

The integrated all-silica-fiber separation and counting device presented above combines elasto-inertial microfluidics with optical fibers and capillaries to open new avenues in ‘Lab-in-a-fiber’ technology for medical diagnostics. This component is capable of sorting micron-sized particles by size and count the separated particles in a single continuous flow. The fiber separation component can be added to a chip-based device as an add-on that may be attractive to the Lab-on-a-chip community.

In this proof-of-principle demonstration of new technique we sorted and analysed ~1400 particles/min. However, the focus of this work is to establish optical fibers and capillaries as an alternative to Lab-on-a-chip technologies that are prevalent in medical diagnostics. At present, the high resistance of the integrated device prevents us from going to higher flow rates at which the throughput of the device can be increased. We believe by optimizing the total component length, increasing the capillary dimension and using a high-pressure pump we can achieve better performance in terms of the throughput. Another important aspect related to the performance of the device is the capability of the separation device to sort particles that have a lower size ratio than what is being demonstrated here. We have used 10 and 1 μm to imitate cells and bacteria. However, it is expected that optimizing the fiber component by changing the separation channel length and flow rates its possible to achieve good separation for the lower size ratio.

In summary, an all-fiber separation component capable of sorting micron-sized particles based on size has been demonstrated. Furthermore, counting of the separated particles in a single-continuous integrated fiber platform is demonstrated.

## Acknowledgement

This work was funded by Knut and Alice Wallenberg foundation.

